# Understanding patterns of abiotic and biotic stress resilience to unleash the potential of crop wild relatives for climate-smart legume breeding

**DOI:** 10.1101/596072

**Authors:** Maarten van Zonneveld, Mohamed Rakha, Shin-yee Tan, Yu-yu Chou, Ching-huan Chang, Joyce Yen, Roland Schafleitner, Ramakrishnan Nair, Ken Naito, Svein Ø. Solberg

## Abstract

Although new varieties are urgently needed for climate-smart legume production, legume breeding lags behind with cereals and underutilizes wild relatives. This paper provides insights in patterns of abiotic and biotic stress resilience of legume crops and wild relatives to enhance the use and conservation of these genetic resources for climate-smart legume breeding. We focus on *Vigna*, a pantropical genus with more than 88 taxa including important crops such as cowpea and mung bean. Sources of pest and disease resistance occur in more than 50 percent of the *Vigna* taxa, which were screened while sources of abiotic stress resilience occur in less than 20 percent of the taxa, which were screened. This difference suggests that *Vigna* taxa co-evolve with pests and diseases while taxa are more conservative to adapt to climatic changes and salinization. Twenty-two *Vigna* taxa are poorly conserved in genebanks or not at all. This germplasm is not available for legume breeding and requires urgent germplasm collecting before these taxa extirpate on farm and in the wild. *Vigna* taxa, which tolerate heat and drought stress are rare compared with taxa, which escape these stresses or tolerate salinity. These rare *Vigna* taxa should be prioritized for conservation and screening for multifunctional traits of combined abiotic and biotic stress resilience. The high presence of salinity tolerance compared with drought stress tolerance, suggests that *Vigna* taxa are good at developing salt-tolerant traits compared with drought-tolerant traits. *Vigna* taxa are therefore of high value for legume production in areas that suffer from salinization.

## Introduction

Legume crops are an important and cheap source of proteins and micronutrients in human diets ^1^. These crops also fix nitrogen because of their symbiosis with *Rhizobium* bacteria, which makes them attractive crops for soil improvement in farming systems ^2^. However, increased abiotic stress such as heat, drought and salinity, and high pressure of disease and insect pests under climate change will decrease yield and quality of existing legume varieties ^3^.

To develop and breed legume varieties that can sustain legume production under global climate change, crop wild relatives and landraces would be important sources for genetic improvement to introgress traits to tolerate, escape, or avoid abiotic stresses and to resist against pests and diseases ^4,5^. So far, genetic improvement of legume crops lags behind compared with cereal crops ^1^, and legume breeders have used little crop wild relatives in developing varieties because of limited information on traits of economic importance in wild species, linkage drag of undesired traits and crossing barriers ^6^. A better understanding of trait diversity in combination with advances in functional genomics and phenotyping will help to fully exploit legume crop diversity to broaden the currently narrow genetic basis in legume breeding ^1,5,6^.

We aim to better understand patterns of abiotic and biotic stress resilience of legume crops and wild relatives for genetic improvement of legume crops focusing on *Vigna*, a complex and pantropical genus of more than 88 taxa. This genus includes a number of important legume crops for food and nutrition security in tropical Asia and Africa such as mung bean (*V. radiata*) and cowpea (*V. unguiculata*) as well as several neglected and underutilized crops such as moth bean (*V. aconitifolia*) and Bambara groundnut (*V. subterranea*).

To provide insights in trait diversity and evolution for *Vigna* introgression breeding, we first examine the distribution of traits related to abiotic and biotic stress resilience across the *Vigna* genus by combining an ecogeographic analysis with a literature review. Second, we develop four *Vigna* gene pools to support introgression breeding by combining existing information on *Vigna* phylogenetics with information on crossing compatibility. Finally, we assess the *ex situ* conservation status of *Vign*a to check out germplasm availability for breeders and to target countries for missions to collect germplasm of *Vigna* taxa, which are not well conserved *in situ* and *ex situ*.

## Methods

Four *Vigna* gene pools with 88 taxa from three subgenera and nine biosystematics sections were defined for nine domesticated *Vigna* taxa as listed in GRIN taxonomy ^7^. Other taxa were excluded because they are part from sections, which are remote to domesticated taxa. We delineated four gene pools following intercrossing studies, which was complemented by genomic, phylogenetic, and biosystematics studies (Text S1). Taxa names were revised according to GRIN taxonomy ^7^ and Iseki et al. ^8^ (Table S1). The *Vigna ex situ* conservation status was determined with data from the 2017 WIEWS database ^9^ and the online platform Genesys ^10^. Breeding objectives were defined on the basis of two key breeding papers of the two most important *Vigna* crops: mung bean and cowpea ^11,12^. These objectives were revised and completed by the coordinator of the International Mungbean Improvement Network (pers. comm., Ramakrishnan Nair, WorldVeg).

### Collection of presence records

A total of 28,313 georeferenced presence records for 84 of the 88 *Vigna* taxa were used for the detection of geographic patterns of taxonomic richness and gaps in genebank collections. The presence records come from four data sources:

1. Presence records from herbaria, which were reported by Tomooka et al. ^13^ were manually curated and where possible georeferenced in Google Earth or with support of www.geonames.org.
2. Presence records from herbaria and living collections, which were stored in the Global Biodiversity Information Facility (https://www.gbif.org/) were collected with the rgbif package ^14^ and manually georeferenced if taxa had less than 30 georeferenced presence records.
3. Presence records from genebanks listed in the 2017 WIEWS database ^9^.
4. Presence records from the *Vigna* genebank collection of the World Vegetable Center (WorldVeg) were manually curated and where possible georeferenced in Google Earth or with support of www.geonames.org.

### Cleaning of presence records

Presence records with inconsistencies between countries as reported in the passport data and at the projected locations outside a border buffer zone of 10 arc minutes were removed following Van Zonneveld et al. ^15^. Coordinates of presence records located in coastal waters within a 10-arc minute buffer zone to the coastline were relocated to the nearest point in the coastline. Presence records with coordinates from country centroids were removed because these points are likely georeferenced at country level with low precision. For each taxon, duplicate records with the same coordinates were removed to reduce sample bias.

Outlier presence records with climate values far beyond taxa’ niche margins were removed from our dataset because these are likely errors in coordinates or taxonomy. We removed outliers when the values of three or more of 19 bioclimatic variables were outside a threshold of 2.5 times the interquartile range below the first quartile or above the third quartile following van Zonneveld et al. ^15^. Climate data were derived from the 2.5-arc minute environmental layers of the 2.0 WorldClim database ^16^.

### Gap analysis

We identified taxonomic and geographic gaps in genebank collections in two steps. First, to target germplasm from taxa and countries underrepresented in *ex situ* collections, we compared sampled taxonomic richness reported by genebanks and living collections with sampled taxonomic richness reported by herbaria. Second, to detect geographic areas where taxa have not been reported before, we applied ecological niche modelling with Maxent, a widely used modelling algorithm to detect areas where climate conditions are suitable for plant species ^17^. We modelled the distribution of each *Vigna* taxon under current climate conditions (1970-2000) using a selection of seven bioclimatic variables available from the WorldClim 2.0 database ^16^ (Text S2): These bioclimatic variables were selected after the removal of correlated variables with a Pearson correlation coefficient of more than 0.7. We used the threshold value of maximum specificity + sensitivity to distinguish suitable from not-suitable areas for taxa presence ^18^. To reduce sampling bias, we first took the average Maxent results for each taxa from three runs each time, using 80% of the randomly resampled records from grid cells with a size corresponding to 10% of the longest inter-point distance after Fourcade et al. ^19^. Second, to allow Maxent to discriminate areas with presence records from the areas with no data, we randomly extracted ten times more background points from the area enclosed by the convex hull polygon based on the presence records, and extended with a buffer corresponding to 10% of the longest inter-point distance. Third, to reduce the risk of including modelled areas where the taxa do not occur in reality, we limited the modelled distribution range by the area enclosed by the convex hull polygon based on the presence records and extended with the buffer around it.

### Assessment of abiotic stress and biotic stress resilience

We did a literature review to assess biotic stress resistance of *Vigna* taxa with a focus on bruchids (*Callosobruchus* spp.), mung bean Yellow Mosaic Disease (YMD), and other pests and diseases, which were included in the breeding objectives. For the assessment of abiotic stress resilience, we carried out an ecogeographic analysis. This analysis was complemented with scores on a 1-5 Likert scale of high to low tolerance of *Vigna* taxa to dehydration and salinity in pot experiments, which were carried out by the Genetic Resources Center of the Japanese Agriculture and Food Science Organization ^8,20^. All taxa, which scored 1 for at least one accession in the pot experiments, were identified in this paper as taxa with a high level of tolerance. Taxa, which scored 2 for at least one accession, were identified as taxa with an intermediate level of tolerance. Taxa with poorer scores were identified as taxa with a low level of tolerance.

The ecogeographic analysis consisted of one-way ANOVAs with ranked climate data to identify *Vigna* taxa, which occur in harsh climate conditions, and which therefore would have acquired traits related to abiotic stress resilience. Climate data was extracted from WorldClim ^16^. *Vigna* taxa were identified for four types of harsh climate conditions:

- *Permanently hot* climate conditions with high annual mean temperature;
- *Seasonally hot* climate conditions with high mean temperatures in the wettest quarter;
- *Permanently dry* climate conditions with low annual rainfall; and
- *Seasonally dry* climate conditions with low rainfall in the wetter quarter.

For *seasonally hot* and *seasonally dry* climate conditions, temperature and rainfall in the wettest quarter were selected because these are critical variables for flowering and seed development of annual plants such as *Vigna* taxa (pers. comm., Ken Naito, Genetic Resources Center, Japanese Agriculture and Food Science Organization). Post-hoc (HSD) Tukey tests were carried out to identify *Vigna* taxa, which occur in harsh climate conditions; these taxa belonged to the HSD group with highest temperature or lowest rainfall values. We identified a second group of *Vigna* taxa, which occur in less harsh climate conditions; these taxa belonged to the HSD group with second-highest temperature or second-lowest rainfall values. Finally, all other *Vigna* taxa belonged to a third group because they occur in less harsh climate conditions. For nine taxa, less than five georeferenced records were available to extract climate data. Because of this lack of sufficient data, these taxa were excluded from the ecogeographic analysis.

### Software

We performed all analyses and graph representations in R version 3.3.3. The code and datasets are available at 10.6084/m9.figshare.7551011 [The link will be made active after acceptance]. Text S3 lists the R packages, which were used.

## Results

### Vigna gene pools

Gene pool A consists of Asian *Vigna* taxa of the subgenus *Ceratotropis*, including mung bean (*V. radiata*), urd bean (*V. mungo*), azuki bean (*V. angularis*), rice bean (*V. umbellata*), creole bean (*V. reflexo-pilosa* var. *glabra*), and moth bean (*V. aconitifolia*). Mung bean is the economically most important *Vigna* of gene pool A (Figure 1). Mung bean can produce mature seeds within two months, which makes it popular for crop diversification of rice and wheat systems ^21^. The domestication centre of mung bean is thought to be in current India. Urd bean is also thought to be domesticated in current India and is genetically close to mung bean, and is especially popular in southern India and Pakistan. Azuki bean is mainly cultivated in Japan and Korea, and is thought to be domesticated in Japan ^22^. Other domesticated taxa are minor crops with important features. Rice bean can grow on poor lands and is mostly cultivated in the tropical Asian highlands ^23^. Creole bean, which is the only known tetraploid *Vigna* taxa ^13^, is cultivated in Vietnam, Philippine, Mauritius, and Tanzania ^13^. Moth bean, another legume crop is thought to be domesticated in current India is cultivated where it is too hot for mung bean cultivation ^23^. In addition, several wild taxa are locally consumed and cultivated in Southeast Asia including jungle bean (*V. trilobata*) and tooapee (*V. trinervia*) ^24,25^.

**Figure 1.**
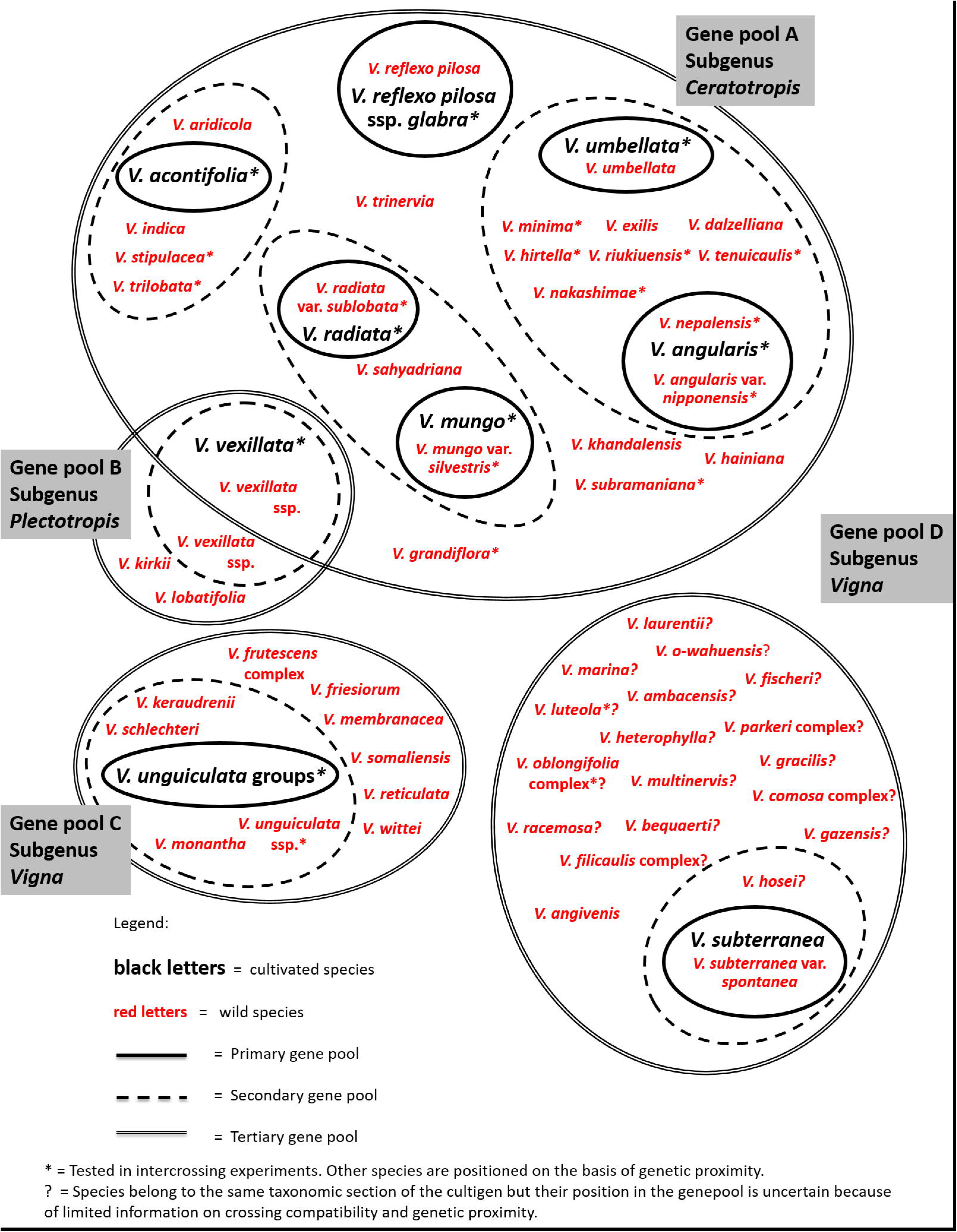
*Vigna* gene pools.

Gene pool B comprises taxa from subgenus *Plectotropis* to which the domesticated tuber cowpea (*V. vexillata*) and its wild relatives belong (Figure 1). There are two types of domesticated tuber cowpea. The tuber type was is domesticated in Asia while the pea type was domesticated in Africa ^26^.

Gene pool C comprises taxa of section *Catiang*, *Macrodontae*, and *Reticulatae* of subgenus *Vigna*. The domesticated taxa in this gene pool are yard-long bean (*V unguiculata* group *sesquipedalis*), which is mostly cultivated in Asia, and cowpea (*V unguiculata* group *unguiculata*), which was domesticated in West and Central Africa. Yard-long bean is different from cowpea by its very long pod (~1 m). People harvest and eat young pods of yard-long bean, rather than the peas.

Finally, gene pool D comprises the domesticated Bambara groundnut (*V. subterranea*) and other taxa of section *Vigna* of subgenus *Vigna* (Figure 1). Bambara groundnut is thought to be domesticated in West Africa ^27^. Farmers cultivate several wild taxa from this gene pool for local consumption, such as *V. marina* and *V. luteola* ^28,29^. Several taxa from this genus are grown widely in the tropics as forages including *V. luteola*, *V. hosei*, and *V. parkeri* ^29^. This pantropical section includes taxa from Africa such as Bambara groundnut, from the Americas such as *V. luteola*, and one species endemic to the pacific: *V. o-wahuensis*. It is the least investigated of all four gene pools. Many taxa of gene pool D have not yet been included in crossing compatibility experiments.

### Breeding objectives

Breeding objectives for mung bean and urd bean related to *abiotic stress* resilience include tolerance to heat stress, water logging, and salinity ^11^. Breeding objectives for cowpea related to *abiotic stress* resilience include drought and heat stress tolerance and phosphorus-use efficiency ^12^.

Important breeding aims related to *biotic stress* for mung bean and urd bean related to disease resistance include YMD, powdery mildew, *Cercospora* leaf spot; and resistance to the following five pests: bruchids, *Thrips* spp., bean flies (*Ophiomyia spp*. and *Melanagromyza* spp.), legume pod borer (*Maruca vitrata*), and whitefly (*Bemisia tabaci*). Breeding objectives for grain cowpea include bacterial blight (*Xanthomonas axonopodis* pv *phaseoli*) and viruses ^12^. Breeding objectives in vegetable cowpea and yard-long bean include resistance against legume pod borer, YMD, and anthracnose (*Colletotrichum destructivum*).

### Abiotic stress resilience

Seven taxa from two gene pools occur in *seasonally hot* climate conditions and nine taxa from three gene pools occur in *seasonally dry* climate conditions (Table 1). In contrast, only two taxa from just one gene pool occur in *permanently hot* climate conditions and two other taxa from two gene pools occur in *permanently dry* climate conditions. Six taxa from three gene pools tolerated high levels of salinity in contrast to only three taxa from two gene pools, which tolerated high levels of dehydration.

**Table 1.**
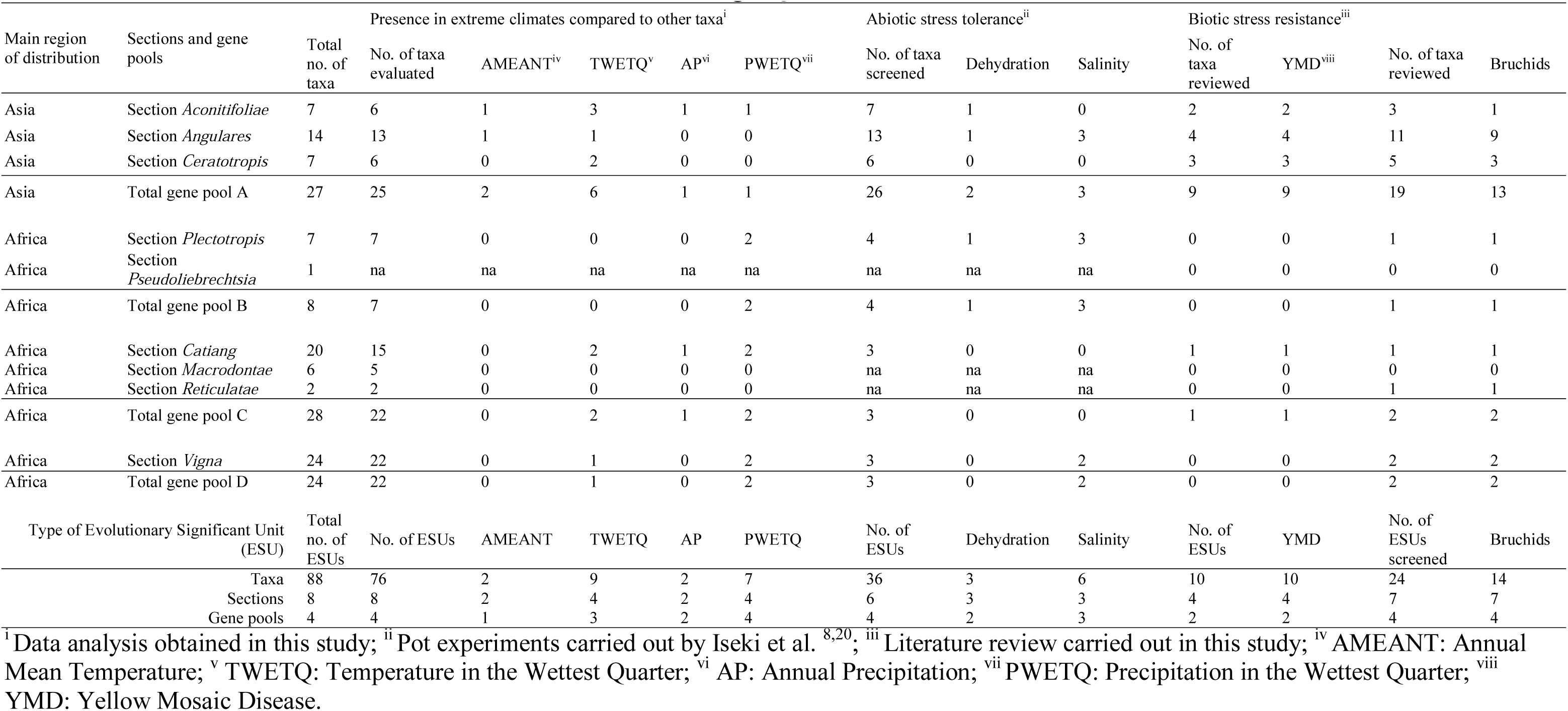
Patterns of abiotic and biotic stress resilience across the *Vigna* genus

The wild relatives *V. trilobata*, *V. vexillata* var. *ovata*, and *V. monantha* and the domesticated moth bean (*V. aconitifolia*) returned highest scores for abiotic stress resilience (Table S2). These four taxa showed high levels of abiotic stress resilience in three of the six variables. *Vigna trilobata* occurs in *seasonally hot* climate conditions and tolerated high levels of dehydration and salinity. *Vigna vexillata* var. *ovata* occurs in *seasonally dry* climate conditions and tolerated high levels of dehydration and salinity. *V. monantha* and *V. aconitifolia* occur in *permanently dry* and *seasonally hot* climate conditions with low rainfall during the wettest quarter. While *V. monantha* was not screened for dehydration tolerance, *V. aconitifolia* accessions did not tolerate high levels of dehydration and salinity.

The wild relatives *V. aridicola*, *V. exilis, V. laurentii* and the domesticated yard-long bean showed high levels of abiotic stress resilience in two of the six variables (Table S2). *Vigna aridicola* occurs in *permanently hot* climate conditions and tolerated high levels of dehydration. *Vigna exilis* occurs in *permanently* and *seasonally hot* climate conditions. This species, however, did not tolerate dehydration or salinity. *Vigna laurentii* occurs *in seasonally hot* and *seasonally dry* climate conditions. The species was not tested for tolerance for dehydration and salinity. Finally, yard-long bean occurs in *seasonally hot* climate conditions and tolerated high levels of salinity.

Thirteen taxa showed high levels of abiotic stress resilience in one of the six variables. *Vigna hainiana*, *V. radiata* var. *sublobata*, and *V. stipulacea* occur in *seasonally hot* climate conditions. *Vigna heterophylla*, *V. kirkii*, and *V. unguiculata* subsp. *stenophylla*, occur in *seasonally dry* climate conditions. Finally, high levels of salinity tolerance were reported for the domesticated cowpea and tuber cowpea, and the wild relatives *V. luteola*, *V. marina, V. nakashimae*, *V. riukiuensis*, and *V. vexillata* var. *macrosperma*.

### Biotic stress resistance

Bruchid resistance was reported in 14 of the 24 taxa (58 %), which were evaluated. These 14 taxa came from all four gene pools (Table S3). However, no resistance against bruchids was reported for mung bean. YMD resistance was reported in all ten taxa (100 %), which were evaluated, and which belonged to gene pool A of mung bean and other Asian *Vigna* crops and gene pool C of cowpea and yard-long bean.

For the other pest and diseases, remarkably less research has been conducted. In gene pool A, only for mung bean and urd bean germplasm, resistance was reported against legume pod borer, whiteflies, stem borer and *Thrips* spp. In this gene pool A, no reports were found for resistance against powdery mildew, bacterial blight, or *Cercospora* leaf spot. In gene pool C, germplasm of the primary gene pool of cow pea and yard-long beam was reported to resist against bacterial blight, *Thrips* spp., and legume pod borer. No reports were found on resistance against anthracnose. Little research was done on biotic stress resistance in gene pools B and D, except for bruchid resistance and cowpea mottle carmovirus (CPMoV).

### *Ex situ* conservation status

In total, 96 institutions conserve *ex situ* 89,288 accessions of the targeted *Vigna* taxa. Eight institutes maintain more than 53,756 of these accessions (60%) (Table S4). As a safety duplicate, 31,500 accessions from 25 taxa are stored in the Svalbard Global Seed Vault (SGSV, 2018, https://www.nordgen.org/sgsv/), mainly from the International Institute of Tropical Agriculture (IITA), the Centro Internacional de Agricultura Tropical (CIAT), and WorldVeg. From the domesticated taxa, IITA holds the largest collections of cowpea and Bambara groundnut. The status of yard-long bean (*V. unguiculata* group *sesquipedalis*) collection is unclear because not all genebanks provided taxonomic data below species level, which is necessary for yard-long bean. WorldVeg holds the largest collection of mung bean and azuki bean. The Indian Bureau of Plant Genetic Resources (NBPGR) holds the largest collections of urd bean, rice bean, and moth bean. CIAT has the largest collection of tuber cowpea. The Genetic Resources Center of the Japanese Agriculture and Food Science Organization has the largest collection of creole bean (*V. reflexo-pilosa*). The Australian Grains genebank and IITA maintain important collections of African wild *Vigna* while the Japanese Agriculture and Food Science Organization and NBPGR hold an important collection of wild Asian *Vigna.* The national botanic garden of Belgium, Meise has the most diverse *Vigna* collection but only keeps a limited number of accessions per taxon.

### Collection gaps

Two Asian *Vigna* and four African *Vigna* are not represented in any of the genebanks, which report to WIEWS: *Vigna sahyadriana, V. indica*, *V. keraudrenii*, *V. monantha*, *V. somaliensis*, and *V. gazensis*. Genebanks maintain less than 10 accessions for nine other Asian *Vigna* and seven other African *Vigna* (Table S4). Priority countries for germplasm collecting missions of these 22 species in Asia are Thailand, India, Sri Lanka, and Myanmar (Table S5). In Africa, priority countries are Madagascar, DRC Congo, South Africa, Benin, Burundi, Somalia, Namibia, and Tanzania. In the Pacific, Hawaii is prioritized to collect *V. o-wahuensis*.

Geographic gaps in Asia with high taxonomic richness reported by herbaria but low coverage reported by genebanks and living collections are Taiwan, Northeast Australia, and India (Figure 2). When comparing the reported overall taxonomic richness with the modelled taxonomic richness, our analysis showed modelled gaps in West Cambodia, Central Thailand, South Vietnam, and coastal India (Figure 2). Geographic gaps in Africa with high reported taxonomic richness but low coverage by genebank collections are Burundi, Benin, and Uganda. In addition, East DRC Congo is a collection gap of modelled richness.

**Figure 2.**
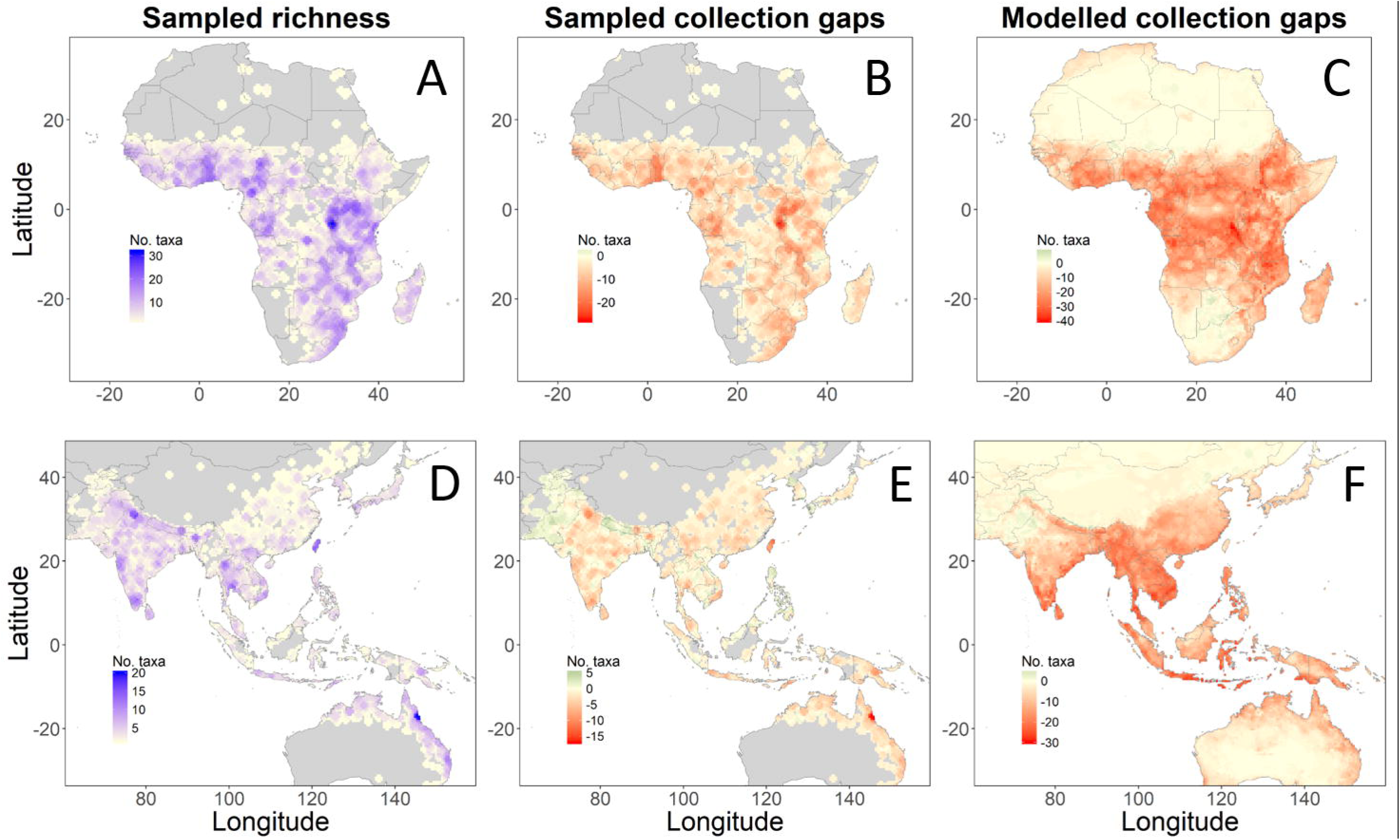
Sampled taxonomic richness and sampled and modelled collection gaps. Panel A and D show sampled taxonomic richness; Panel B and E show sampled taxonomic richness, which is not conserved *ex situ*; Panel C and F show gaps where a high number of taxa are modelled to occur but are not reported in herbaria, living collections, and genebanks.

## Discussion

Our findings show that sources of pest and disease resistance occur in more than 50 percent of the taxa, which were screened. In contrast, sources of abiotic stress resilience occur in less than 20 percent of the taxa, which were screened. We therefore hypothesize that during evolution, *Vigna* taxa have been conservative in acquiring traits related to abiotic stress resilience compared with pest and diseases resistance. This hypothesis can be tested by screening of gene pools of other legume crops. We only found one study, which simultaneously screened legume wild relatives for abiotic stress and biotic stress resilience ^30^. This study with eight chick pea wild relatives (*Cicer* spp.) focuses on cold stress rather than heat stress in combination with biotic stress resistance. Nevertheless, the study confirms our hypothesis because the researchers identified only two wild *Cicer* species tolerating cold stress compared with four or more *Cicer* species resisting bruchids and other pests and diseases.

Even though some traits, which cope with abiotic stress, such as waterlogging, are regulated by just one or few genes ^31,32^, several traits related to drought and heat stress tolerance could be difficult for plants to acquire because they are related to numerous genes ^32^ or because of trade-offs with other traits. This could explain the low percentage of *Vigna*, which tolerate dehydration, and of *Vigna*, which occur in *permanently hot* and *dry* climate conditions.

Nine percent of the taxa screened in the ecogeographic analysis occur in *seasonally hot* or *seasonally dry* climate conditions, where it is possible to escape heat and drought stress rather than to tolerate heat and drought stress. In contrast, the three percent of the taxa, which occur in *permanently hot* and *permanently dry* climate conditions require traits to tolerate continuous heat and drought stress. This finding suggests that during evolution *Vigna* taxa more easily acquire phenological traits for short life cycles to escape drought and heat stresses compared with acquiring physiological traits to tolerate these stresses continuously.

Resistance against pest and diseases has often been a result from continuous and recent co-evolution in the geographic areas of occurrence and pressure of pest and diseases ^33^. Pest pressure can change genotype frequencies in a plant population within just a few generations ^34,35^. For wild tomato relatives, the density of trichomes and levels of acylsugar concentrations, related to direct pest defense, correspond with pest pressure and the geographic distribution of these pests ^36,37^. Co-evolution can explain the high percentage of taxa, which possesses biotic stress resistance compared with heat and drought stress resilience.

The presence of salinity tolerance in 17 percent of taxa compared with three percent drought-tolerant taxa, suggest that *Vigna* taxa are good at developing salt-tolerant traits compared with drought-tolerant traits, and which would only require a few gene changes ^8^. Many wild *Vigna* taxa occur naturally in coastal areas and are adapted to a saline environment ^28^. This makes *Vigna* genetic resources of high value for legume production in areas, which suffer from salinization.

Our findings suggest that the section *Aconitifolia* has high levels of heat and drought stress resilience. This section includes moth bean, which occurs in *permanently dry* and *seasonally hot* climate conditions. This finding contrasts with moth bean`s low dehydration tolerance ^20^. This discrepancy could be explained by several reasons. First, the pot experiment measured dehydration tolerance at seedling stage while plants have also acquired traits to tolerate or escape drought stress during the vegetative or reproductive stage, which are captured in the ecogeographic analysis with presence records of wild and cultivated populations of plants, which complete their life cycle. Second, the limited number of accessions of moth bean in the pot experiments may not reflect the full intra-specific genetic variation of this species. Finally, *Vigna* taxa, such as moth bean may develop a third type of resilience besides *tolerating* or *escaping* drought stress; they could *avoid* drought stress, for example by developing a large root/shoot ratio such as several tuber *Vigna* taxa do as storage organs for dry months.

Vice versa, *V. aridicola* from section *Aconitifoliae* tolerated drought dehydration while the ecogeographic analysis indicated that this species occurs in humid regions. It could be that our analysis does not reflect the complete climatic ranges because we only were able to collect 14 records of this species.

Cowpea and mung bean are the two economically most important *Vigna* crops. However, only few studies were found on biotic stress resilience of cowpea wild relatives compared with mung bean wild relatives. It could be that cowpea breeders do not yet need to use cowpea wild relatives for biotic stress resilience because the many botanical varieties in cowpea`s primary gene pool provide sufficient variation for finding traits. Another possibility could be that cowpea is difficult to cross with close relatives compared with mung bean and its close relatives, which would need to be tested in future studies.

We developed four *Vigna* gene pools. Even though, these gene pools may be modified after new studies on crossing compatibility, they provide a helpful structure to examine trait distribution and to support introgressions in *Vigna* breeding. For many *Vigna* taxa, genetic relationships and crossing compatibility are still poorly understood. This is especially true for the pantropical *Vigna* section of subsgenus *Vigna*, which includes taxa from Africa, the Americas, and the Pacific. Domestication and origin of *Vigna* taxa is largely unknown. For example, where did the *Vigna* genus originate and how did the genus spread across Africa, Asia, the Americas, and the Pacific?

Twenty-six percent of Vigna species requires urgent germplasm collecting efforts because they are not- or under-represented in genebanks. Two Asian *Vigna* species and four African *Vigna* species require urgent efforts of germplasm collecting because these species are not reported in any genebank. Nine other Asian *Vigna* species and seven other African *Vigna* species also require urgent efforts of germplasm collecting because genebanks maintain less than 10 accessions of these species. Targeted Asian countries for germplasm collecting efforts include India, Thailand, Thailand, Myanmar, Australia, and Taiwan. In Africa, our sampled and modelled richness analyses indicate Burundi, DRC Congo, and Madagascar as priority countries for *Vigna* germplasm collecting efforts.

*Vigna* taxa with high levels of heat and drought stress tolerance are rare. We therefore propose to use the presence of these type of traits as a criterion to prioritize taxa for conservation. The section Aconitifoliae requires urgent conservation efforts when considering this criterion in combination with poor genebank coverage. Five out of the seven *Aconitifoliae* taxa are poorly conserved *ex situ* with 10 or less accessions in genebanks while this section includes several traits related to abiotic stress tolerance. Taxa from this section mainly occur in India and Sri Lanka, which are priority countries for germplasm collecting. *Vigna exilis*, which occurs in *permanently hot* climate conditions and V*. monantha*, which occurs in *permanently dry* conditions, also require urgent conservention because these species are not represented in genebanks, except for one accession of *V. exilis*.

*Vigna monantha* and another African *Vigna*, *V. keraudrenii* require urgent *in situ* and *ex situ* conservation efforts because these species are endangered according to the IUCN Red List ^38^. Until now, the IUCN has evaluated only six *Vigna* species as part of the Red List. The inclusion of more *Vigna* taxa on the IUCN Red List will help to better understand their *in situ* conservation status.

The pacific *V. o-wahuensis*, endemic to Hawaii, is critically endangered according to the U.S. Endangered Species Act ^39^. Fortunately, 27 accessions have been safeguarded in the Lyon Arboretum, Hawaiian Rare Plant Program (pers. comm. Marian Chau, Lyon Arboretum). Strengthening collaboration between genebanks and botanical gardens such as the Lyon Arboretum and Meuse Belgium, would further enhance *ex situ* conservation and germplasm availability of *Vigna*.

To date screening and understanding of the ability of *Vigna* taxa to adapt against combined stresses is largely unknown. These taxa would be priority for screening for introgressions in climate-smart legume breeding. These taxa could have developed independent traits to cope with abiotic and biotic stress or multifunctional traits, such as high levels of antioxidant capacity, which can cope with both abiotic and biotic stress at the same time ^40^. With functional genomics, saturated markers could help to determine the precise position and size of wild introgressions to introduce genes related to these desired traits and to minimize linkage drag of undesired traits.

Our analysis suggests that many taxa from the section *Aconitifoliae* show high levels of abiotic stress resilience, but only a limited number of accessions from this section were evaluated for resistance against pests and diseases. *Vigna* taxa with high levels of abiotic stress resilience require further evaluation under different abiotic and biotic stress combinations.

We propose trichomes as a promising multifunctional trait in *Vigna* taxa for further screening because this trichomes help plants to cope with both abiotic and biotic stress ^41^. High density of trichomes on the pods of mung bean and cowpea complicates the mobility of the adult of bruchids and pod borers over the pods and decrease pest infestations ^42,43^. Numerous studies show that glandular trichomes in tomato wild relatives produce secondary metabolites including acylsugars, methylketones, and sesquiterpenes, which intoxicate, repel, or trap pests ^44,45^. At the same time, trichomes may increase abiotic stress tolerance by reducing leaf radiation absorbance and facilitating condensation of air moisture onto the plant surface, among other functions ^46^. Further research will reveal the relationships between trichomes types and densities, trichome evolution, and abiotic and biotic stress resilience in *Vigna* taxa.

## Acknowledgements

Funding for the World Vegetable Center’s general research activities is provided by core donors: Republic of China (Taiwan), UK aid from the UK government, United States Agency for International Development (USAID), Australian Centre for International Agricultural Research (ACIAR), the Federal Ministry for Economic Cooperation and Development of Germany, Thailand, Philippines, Korea, and Japan.

## Contributions

M.vZ., M.R., Y.C., and S.Ø.S. conceived the ideas; M.vZ., M.R., S.T., R.N., C.C., J.Y., K.N., and S.Ø.S. contributed to data and information collection; M.vZ. and S.T. analysed the data; M.vZ. and M.R. wrote the paper; R.S. provided ideas and critique in the analysis and writing process.

## Additional information

The authors declare no competing interests.

