## Supplementary information for "Understanding patterns of abiotic and biotic stress resilience to unleash the potential of crop wild relatives for climate-smart legume breeding"

**This document includes the following supplementary texts, tables and references:**

**Text S1.** Literature review of crossing compatibility and phylogenetic analysis to develop four gene pools

**Text S2.** Selected bioclimatic variables for Maxent distribution modelling

**Text S3.** R packages used

**Table S1.** Accepted *Vigna* names and taxonomic classification following GRIN taxonomy ^1^ and Iseki et al. ^2^

**Table S2.** Abiotic stress resistance of *Vigna* species

**Table S3.** Biotic stress tolerance of *Vigna* species

**Table S4.** Reported accessions conserved *ex situ* in 2017

**Table S5.** Targeted countries for collecting of *V.* species with less than 10 genebank accessions

**References**

**Text S1.** Literature review of crossing compatibility and phylogenetic analysis to develop four gene pools

**Domesticated *Vigna* taxa**

**Gene pool A. Subgenus *Ceratotropis*:** *Vigna aconitifolia* ^3,4^, *V. angularis* ^3,4^, *V. mungo* ^3,4^, *V. radiata*^3,4^, *V. reflexo-pilosa* var. *glabra* ^3,4^, *V. umbellata*^3,4^.

**Gene pool B. Subgenus *Plectotropis*:** *V. vexillata* ^3,4^.

**Gene pool C. Subgenus *Vigna*:** *V. unguiculata* goups ^4^

**Gene pool D. subgenus *Vigna*:** *V. subterranea* ^4^.

**Gene pool A. Subgenus *Ceratotropis***

*Vigna angularis* x *V. hirtella:* secondary gene pool. Fertile F1 hybrids were produced between male *V. angularis* and female *V. hirtella* ^5,6^.

*Vigna angularis* x *V. minima*, *V. nakashimae*, *V. riukiuensis, V. tenuicaulis*: secondary gene pool. Fertile F1 hybrids were developed in all cases ^6^.

*Vigna angularis* x *V. nepalensis*: primary gene pool. Fertile F1 hybrids were developed ^6^.

*Vigna angularis* x *V. umbellata*: secondary gene pool. Only crosses with female *V. umbellata* plants produces fertile F1 hybrids ^6,7^.

*Vigna radiata* x *V. mungo:* secondary gene pool. Hybrids between *V. radiata* and *V. mungo* can be produced when *V. radiata* is the female parent. F1 seeds are often shrivelled and do not germinate. F1 plants show low fertility^6,8^.

*Vigna radiata* x *V. radiata* var. *sublobata:* primary gene pool. Crosses produce fertile F1, although in some crosses there is reduced F1 pollen sustainability ^6,9^.

*Vigna radiata* x *V. trilobata:* tertiary gene pool. Crosses with female and male *V. radiata* plants return low pod set and low germination rates ^10^. Successful F1 hybrids were made (person communication Ram Nair, World Vegetable Center). Moderate crossability was observed between *V. radiata* and *V. trilobata* (8.48%) ^11^.

*Vigna radiata* x *V. vexillata:* tertiary gene pool*.* Crosses with female and male *V. radiata* plants return low pod set and low germination rates ^10^.

*Vigna radiata* x *V. grandiflora:* tertiary gene pool. Embryo rescue was necessary to obtain F1 hybrids ^6,12^

*Vigna radiata* and *V. mungo* x *V. subramaniana:* tertiary gene pool. Embryo rescue was necessary to obtain F1 hybrids ^5,6^.

*Vigna radiata* x *V. aconitifolia*: tertiary gene pool. Moderate crossability was observed between female *V. radiata* and male *V. trilobata* with good seed production (8.48%) ^11^.

*Vigna radiata* x *V. stipulacea*: tertiary gene pool. Low podset between female V. radiata and male *V. stipulacea* ^10^*.*

*V. radiata x Vigna umbellata*: tertiary gene pool. Sterile F1 hybrids have been produced by crossing female *V. radiata* and male *V. umbellata* ^6,13^.

*Vigna radiata* x *V. reflexo-pilosa* var. *glabra:* tertiary gene pool. Crosses between male *V. radiata* and female *V. reflexo-pilosa* var. *glabra* returned sterile F1 hybrids ^6,14^.

*Vigna mungo x V. mungo var. silvestris:* primary gene pool. Crosses produce fertile F1, although in some crosses there is reduced F1 pollen sustainability ^6,9^.

*Vigna umbellata* x *V. minima*, *V. nakashimae*, *V. riukiuensis, V. nepalensis, V. tenuicaulis:* secondary gene pool. Only crosses with female *V. umbellata* plants produces fertile F1 hybrids ^6^.

*Vigna umbellata* x *V. minima:* tertiary gene pool. Crosses produce sterile F1. V. minima is considered as the tertiary gene pool of the rice bean ^6,15^.

*Vigna minima* x *V. hirtella*. Primary gene pool. Natural hybridization results in prima fertile F1 ^3^.

**Gene pool B. Subgenus *Plectotropis***

*Vigna vexillata* x *Vigna vexillata* var. *davyi:* secondary gene pool. Crosses between pairs are partially fertile ^16^.

*Vigna vexillata* x *Vigna vexillata* var. *angustifolia:* secondary gene pool. Pollen fertility between 47% ^16^.

*Vigna vexillata* x *Vigna vexillata* var. *vexillata:* secondary gene pool. Pollen fertility between 59% ^16^.

*Vigna vexillata* x *Vigna unguiculata:* seperate gene pools. No viable hybrids even not after embryo rescue ^17^.

**Gene pool C. Subgenus *Vigna***

*Vigna schlechteri* and *V. keraudrenii* are part of the secondary gene pool of *V. unguiculata* ^18^*.*

*Vigna unguiculata* x *Vigna unguiculata* ssp. *stenophylla:* secondary gene pool. The F1 hybrids of crossings between these two species were partially fertile ^16^.

*Vigna unguiculata* x *Vigna unguiculata* ssp. d*ekindtiana:* tertiary gene pool. A successful cross was made with the help of embryo rescue ^16^.

**Gene pool D. subgenus *Vigna***

*Vigna oblongifolia* x *V. luteola:* tertiary gene pool. A successful cross was made with the help of embryo rescue ^16^.

**Additional species were added to the gene pool model based on taxonomic and phylogenetic analysis:**

**Gene pool A. *Ceratotropis*:** *Vigna dalzelliana* ^2^; *V. hainiana* ^19^; *Vigna indica* ^2^; *V. khandalensis* ^2^; *V. nepalensis* ^19,20^; *V. sahyadriana* ^2^; *V. stipulacea* ^19^; and *V. trinervia* ^2,19^*.*

**Gene pool B. *Plectotropics*:** *Vigna angivensis* ^21^; *V. lobatifolia* ^22^; and *V. vexilliata* spp. ^2,19,21^*.*

**Gene pool C. *Vigna*:** *Vigna friesiorum* ^23^; *V. frutescens* ssp ^21,23^; *V. keraudrenii* ^4^; *V. reticulata* ^21^; and *V. schlechteri* ^4,18^*.*

**Gene pool D. *Vigna*:** *Vigna ambacensis* ^21^; *V. fischeri* ^23^; *V. gracilis* ^21^; *V. heterophylla* ^21^; *V. hosei* ^4,21^; *V. laurentii* ^21^; *Vigna luteola* ^2,21,24^; *Vigna marina* ^2,21,24^; *V. membranacea* ^21,23^; *V. multinervis* ^21,25^; *V. parkeri* ssp*.* ^21^; *V. racemosa* ^21,25^; and *V. subterranea* ^2,24^*.*

**Text S2.** Selected bioclimatic variables for Maxent distribution modelling

- Mean Diurnal Range (Mean of monthly (max temp - min temp));
- Mean Temperature of Warmest Quarter;
- Mean Temperature of Coldest Quarter;
- Annual Precipitation;
- Precipitation of Driest Month;
- Precipitation Seasonality (Coefficient of Variation);
- Precipitation of Warmest Quarter;
- Precipitation of Coldest Quarter.

**Text S3.** R packages used

The following R packages were used in the collection of presence records, Maxent distribution modelling, and the ecogeographic analysis: rgbif´ ^26^, `raster´^27^, `dismo´ ^28^, `sp´ ^29^, `rgeos´ ^30^, `rgdal´ ^31^, `geosphere´^32^ , and `maptools´ ^33^, `ggplot´ ^34^, `doBy´ ^35^, and `agricolae´ ^36^.

**Table S1.** Accepted *Vigna* names and taxonomic classification following GRIN taxonomy ^1^ and Iseki et al. ^2^

| Subgenus | Section | Species | Complete scientific name | Taxonomy |
| --- | --- | --- | --- | --- |
| *Ceratotropis* | *Angulares* | *V. angularis* | *V. angularis* (Willd.) Ohwi & H. Ohashi | ^1^ |
| *Ceratotropis* | *Angulares* | *V. angularis* var. *nipponensis* | *V. angularis* var. *nipponensis* (Ohwi) Ohwi & H. Ohashi | ^1^ |
| *Ceratotropis* | *Angulares* | *V. dalzelliana* | *V. dalzelliana* (Kuntze) Verdc. | ^1^ |
| *Ceratotropis* | *Angulares* | *V. exilis* | *V. exilis* Tateishi & Maxted | ^1^ |
| *Ceratotropis* | *Angulares* | *V. hirtella* | *V. hirtella* Ridl. | ^1^ |
| *Ceratotropis* | *Angulares* | *V. minima* | *V. minima* (Roxb.) Ohwi & H. Ohashi | ^1^ |
| *Ceratotropis* | *Angulares* | *V. nakashimae* | *V. nakashimae* (Ohwi) Ohwi & H. Ohashi | ^1^ |
| *Ceratotropis* | *Angulares* | *V. nepalensis* | *V. nepalensis* Tateishi & Maxted | ^1^ |
| *Ceratotropis* | *Angulares* | *V. reflexopilosa* | *V. reflexopilosa* Hayata | ^1^ |
| *Ceratotropis* | *Angulares* | *V. reflexopilosa* subsp. *glabra* | *V. reflexopilosa* subsp. glabra (Roxb.) N. Tomooka & Maxted | ^1^ |
| *Ceratotropis* | *Angulares* | *V. riukiuensis* | *V. riukiuensis* (Ohwi) Ohwi & H. Ohashi | ^1^ |
| *Ceratotropis* | *Angulares* | *V. tenuicaulis* | *V. tenuicaulis* N. Tomooka & Maxted | ^1^ |
| *Ceratotropis* | *Angulares* | *V. trinervia* | *V. trinervia* (B. Heyne ex Wight & Arn.) Tateishi & Maxted | ^1^ |
| *Ceratotropis* | *Angulares* | *V. umbellata* | *V. umbellata* (Thunb.) Ohwi & H. Ohashi | ^1^ |
| *Ceratotropis* | *Ceratotropis* | *V. grandiflora* | *V. grandiflora* (Prain) Tateishi & Maxted | ^1^ |
| *Ceratotropis* | *Ceratotropis* | *V. hainiana* |  | ^2^ |
| *Ceratotropis* | *Ceratotropis* | *V. mungo* | *V. mungo* (L.) Hepper | ^1^ |
| *Ceratotropis* | *Ceratotropis* | *V. mungo* var. *silvestris* | *V. mungo* var. *silvestris* Lukoki et al. | ^1^ |
| *Ceratotropis* | *Ceratotropis* | *V. radiata* | *V. radiata* (L.) R. Wilczek | ^1^ |
| *Ceratotropis* | *Ceratotropis* | *V. radiata* var. *sublobata* | *V. radiata* var. *sublobata* (Roxb.) Verdc. | ^1^ |
| *Ceratotropis* | *Ceratotropis* | *V. sahyadriana* |  | ^2^ |
| *Ceratotropis* | *Aconitifoliae* | *V. aconitifolia* | *V. aconitifolia* (Jacq.) Maréchal | ^1^ |
| *Ceratotropis* | *Aconitifoliae* | *V. aridicola* | *V. aridicola* N. Tomooka & Maxted | ^1^ |
| *Ceratotropis* | *Aconitifoliae* | *V. indica* |  | ^2^ |
| *Ceratotropis* | *Aconitifoliae* | *V. khandalensis* | *V. khandalensis* (Santapau) Sundararagh. & Wadhwa | ^1^ |
| *Ceratotropis* | *Aconitifoliae* | *V. stipulacea* | *V. stipulacea* (Lam.) Kuntze | ^1^ |
| *Ceratotropis* | *Aconitifoliae* | *V. subramaniana* | *V. subramaniana* (Babu ex Raizada) Raizada | ^1^ |
| *Ceratotropis* | *Aconitifoliae* | *V. trilobata* | *V. trilobata* (L.) Verdc. | ^1^ |
| *Plectrotropis* | *Plectotropis* | *V. kirkii* | *V. kirkii* (Baker) J. B. Gillett | ^1^ |
| *Plectrotropis* | *Plectotropis* | *V. vexillata* | *V. vexillata* (L.) A. Rich. | ^1^ |
| *Plectrotropis* | *Plectotropis* | *V. vexillata* var. *angustifolia* | *V. vexillata* var. *angustifolia* (Schumach.) Baker | ^1^ |
| *Plectrotropis* | *Plectotropis* | *V. vexillata* var. *davyi* | *V. vexillata* var. *davyi* (Bolus) B. J. Pienaar | ^1^ |
| *Plectrotropis* | *Plectotropis* | *V. vexillata* var. *macrosperma* | *V. vexillata* var. *macrosperma* Maréchal et al. | ^1^ |
| *Plectrotropis* | *Plectotropis* | *V. vexillata* var. *ovata* | *V. vexillata* var. *ovata* (E. Mey.) B. J. Pienaar, nom. inval. | ^1^ |
| *Plectrotropis* | *Plectotropis* | *V. vexillata* var. *vexillata* | *V. vexillata* var. *vexillat*a | ^1^ |
| *Plectrotropis* | *Plectotropis* | *V. vexillata* var. *youngiana* | *V. vexillata* var. *youngiana* F. M. Bailey | ^1^ |
| *Plectrotropis* | *Pseudoliebrechtsia* | *V. lobatifolia* | *V. lobatifolia* Baker | ^1^ |
| *Vigna* | *Catiang* | *V. keraudrenii* | *V. keraudrenii* Du Puy & Labat | ^1^ |
| *Vigna* | *Catiang* | *V. monantha* | *V. monantha* Thulin | ^1^ |
| *Vigna* | *Catiang* | *V. schlechteri* | *V. schlechteri* Harms | ^1^ |
| *Vigna* | *Catiang* | *V. unguiculata* | *V. unguiculata* (L.) Walp. | ^1^ |
| *Vigna* | *Catiang* | *V. unguiculata* group *biflora* | *V. unguiculata* (L.) Walp. group *biflora* | ^1^ |
| *Vigna* | *Catiang* | *V. unguiculata* subsp. *baoulensis* | *V. unguiculata* (L.) Walp. subsp. *baoulensis* (A. Chev.) Pasquet | ^1^ |
| *Vigna* | *Catiang* | *V. unguiculata* subsp. *protracta* | *V. unguiculata* (L.) Walp. subsp. *protracta* (E. Mey.) B. J. Pienaar | ^1^ |
| *Vigna* | *Catiang* | *V. unguiculata* group *melanophthalmus* | *V. unguiculata* (L.) Walp. group *melanophthalmu*s | ^1^ |
| *Vigna* | *Catiang* | *V. unguiculata* group *unguiculata* | *V. unguiculata* (L.) Walp. group *unguiculata* | ^1^ |
| *Vigna* | *Catiang* | *V. unguiculata* subsp. *aduensis* | *V. unguiculata* subsp. *aduensis* Pasquet | ^1^ |
| *Vigna* | *Catiang* | *V. unguiculata* subsp. *burundiensis* | *V. unguiculata* (L.) Walp. subsp. *burundiensis* Pasquet | ^1^ |
| *Vigna* | *Catiang* | *V. unguiculata subsp. letouzeyi* | *V. unguiculata* (L.) Walp. subsp. *letouzeyi* Pasquet | ^1^ |
| *Vigna* | *Catiang* | *V. unguiculata subsp. pawekiae* | *V. unguiculata* (L.) Walp. subsp. *pawekiae* Pasquet | ^1^ |
| *Vigna* | *Catiang* | *V. unguiculata* subsp. *tenuis* | *V. unguiculata* subsp. *tenuis* (E. Mey.) Maréchal et al. | ^1^ |
| *Vigna* | *Catiang* | *V. unguiculata* subsp. *unguiculata* | *V. unguiculata* subsp. *unguiculata* | ^1^ |
| *Vigna* | *Catiang* | *V. unguiculata* subsp. *stenophylla* | *V. unguiculata* subsp. *stenophylla* (Harv.) Maréchal et al. | ^1^ |
| *Vigna* | *Catiang* | *V. unguiculata* var*. spontanea* | *V. unguiculata* var. *spontanea* (Schweinf.) Pasquet | ^1^ |
| *Vigna* | *Catiang* | *V. unguiculata* group *sesquipedalis* | *V. unguiculata* group *sesquipedalis* | ^1^ |
| *Vigna* | *Catiang* | *V. unguiculata* subsp. *alba* | *V. unguiculata* subsp*. alba* (G. Don) Pasquet | ^1^ |
| *Vigna* | *Catiang* | *V. unguiculata* subsp. *dekindtiana* | *V. unguiculata* subsp. *dekindtiana* (Harms) Verdc. | ^1^ |
| *Vigna* | *Catiang* | *V. unguiculata* subsp*. pubescens* | *V. unguiculata* subsp. *pubescens* (R. Wilczek) Pasquet | ^1^ |
| *Vigna* | *Macrodontae* | *V. friesiorum* | *V. friesiorum* Harms | ^1^ |
| *Vigna* | *Macrodontae* | *V. frutescens* | *V. frutescens* A. Rich. | ^1^ |
| *Vigna* | *Macrodontae* | *V. frutescens* subsp. *incana* | *V. frutescens* subsp. *incana* (Taub.) Verdc. | ^1^ |
| *Vigna* | *Macrodontae* | *V. frutescens* var. *buchneri* | *V. frutescens* var. *buchneri* (Harms) Verdc. | ^1^ |
| *Vigna* | *Macrodontae* | *V. membranacea* | *V. membranacea* A. Rich. | ^1^ |
| *Vigna* | *Macrodontae* | *V. somaliensis* | *V. somaliensis* Baker f. | ^1^ |
| *Vigna* | *Reticulatae* | *V. reticulate* | *V. reticulata* Hook. f. | ^1^ |
| *Vigna* | *Reticulatae* | *V. wittei* | *V. wittei* Baker f. | ^1^ |
| *Vigna* | *Vigna* | *V. ambacensis* | *V. ambacensis* Welw. ex Baker | ^1^ |
| *Vigna* | *Vigna* | *V. angivensis* | *V. angivensis* Baker | ^1^ |
| *Vigna* | *Vigna* | *V. bequaertii* | *V. bequaertii* R. Wilczek | ^1^ |
| *Vigna* | *Vigna* | *V. comosa* | *V. comosa* Baker | ^1^ |
| *Vigna* | *Vigna* | *V. comosa* var. *lebrunii* | *V. comosa* var. *lebruni*i | ^1^ |
| *Vigna* | *Vigna* | *V. filicaulis* | *V. filicaulis* Hepper | ^1^ |
| *Vigna* | *Vigna* | *V. filicaulis* var*. pseudovenulosa* | *V. filicaulis* var. *pseudovenulosa* Maréchal et al. | ^1^ |
| *Vigna* | *Vigna* | *V. fischeri* | *V. fischeri* Harms | ^1^ |
| *Vigna* | *Vigna* | *V. gazensis* | *V. gazensis* Baker f. | ^1^ |
| *Vigna* | *Vigna* | *V. gracilis* | *V. gracilis* (Guill. & Perr.) Hook. f. | ^1^ |
| *Vigna* | *Vigna* | *V. heterophylla* | *V. heterophylla* A. Rich. | ^1^ |
| *Vigna* | *Vigna* | *V. hosei* | *V. hosei* (Craib) Backer | ^1^ |
| *Vigna* | *Vigna* | *V. laurentii* | *V. laurentii* De Wild. | ^1^ |
| *Vigna* | *Vigna* | *V. luteola* | *V. luteola* (Jacq.) Benth. | ^1^ |
| *Vigna* | *Vigna* | *V. marina* | *V. marina* (Burm.) Merr. | ^1^ |
| *Vigna* | *Vigna* | *V. multinervis* | *V. multinervis* Hutch. & Dalziel | ^1^ |
| *Vigna* | *Vigna* | *V. oblongifolia* | *V. oblongifolia* A. Rich. | ^1^ |
| *Vigna* | *Vigna* | *V. o-wahuensis* | *V. o-wahuensis* Vogel | ^1^ |
| *Vigna* | *Vigna* | *V. parkeri* | *V. parkeri* Baker | ^1^ |
| *Vigna* | *Vigna* | *V. parkeri* subsp. *acutifolia* | *V. parkeri* subsp*. acutifolia* Verdc. | ^1^ |
| *Vigna* | *Vigna* | *V. parkeri* subsp. *maranguensis* | *V. parkeri* subsp. *maranguensi*s (Taub.) Verdc. | ^1^ |
| *Vigna* | *Vigna* | *V. racemosa* | *V. racemosa* (G. Don) Hutch. & Dalziel | ^1^ |
| *Vigna* | *Vigna* | *V. subterranea* | *V. subterranea* (L.) Verdc. | ^1^ |
| *Vigna* | *Vigna* | *V. subterranea* var. *spontanea* | *V. subterranea* var. *spontanea* (Harms) Pasquet | ^1^ |
| *Vigna* | *Vigna* | *V. oblongifolia* var. *parviflora* | *V. oblongifolia* A. Rich. var. *parviflora* (Welw. ex Baker) Verdc. | ^1^ |

**Table S2.** Abiotic stress tolerance of *Vigna* species

|  |  |  |  |  | **Heat** | | **Drought** | | | **Salt** |
| --- | --- | --- | --- | --- | --- | --- | --- | --- | --- | --- |
| Gene  pool | Subgenus | Section | Species | no. of records | Mean AMEANT (°C)^i^ | Mean TWETQ (°C)^i ii^ | Mean AP (mm)^iii^ | Mean PWETQ (mm)^iv^ | Tolerance score^v^ | Tolerance score^vi^ |
| A | *Ceratotropis* | *Aconitifoliae* | *V. aconitifolia* | 172 | 23.7 | 27.4 | 623 | 406 | 2,2,2,3,4,4 | 3,3,3,4,4,4 |
| A | *Ceratotropis* | *Aconitifoliae* | *V. aridicola* | 14 | 27.5 | 26.1 | 1963 | 941 | 1,3,3 | 3,4,4 |
| A | *Ceratotropis* | *Aconitifoliae* | *V. indica* | 0 | na | na | na | na | 2 | 3 |
| A | *Ceratotropis* | *Aconitifoliae* | *V. khandalensis* | 49 | 23.4 | 23.4 | 1788 | 1350 | 4 | 4 |
| A | *Ceratotropis* | *Aconitifoliae* | *V. stipulacea* | 28 | 26.5 | 26.3 | 1367 | 724 | 4,4 | 2,3 |
| A | *Ceratotropis* | *Aconitifoliae* | *V. subramaniana* | 13 | 23.3 | 25.1 | 1252 | 820 | 4,5 | 3,4 |
| A | *Ceratotropis* | *Aconitifoliae* | *V. trilobata* | 267 | 26.5 | 27.0 | 1163 | 704 | 1,3,4 | 1,1,3 |
| A | *Ceratotropis* | *Angulares* | *V. angularis* | 1033 | 13.1 | 21.2 | 1464 | 588 | na | na |
| A | *Ceratotropis* | *Angulares* | *V. angularis* var. *nipponensis* | 303 | 14.2 | 22.1 | 1680 | 663 | 5,5 | 4,4 |
| A | *Ceratotropis* | *Angulares* | *V. dalzelliana* | 106 | 24.1 | 24.6 | 1822 | 1053 | 4,4 | 2,3 |
| A | *Ceratotropis* | *Angulares* | *V. exilis* | 17 | 27.6 | 27.5 | 1266 | 644 | 4 | 2 |
| A | *Ceratotropis* | *Angulares* | *V. hirtella* | 32 | 23.0 | 24.6 | 1750 | 955 | 4,5,5,5 | 3,4,4,4 |
| A | *Ceratotropis* | *Angulares* | *V. minima* | 303 | 22.6 | 25.2 | 2073 | 949 | 3,3,3 | 2,2,3 |
| A | *Ceratotropis* | *Angulares* | *V. nakashimae* | 38 | 14.4 | 22.8 | 1712 | 754 | 4 | 1 |
| A | *Ceratotropis* | *Angulares* | *V. nepalensis* | 11 | 15.7 | 20.0 | 2248 | 1374 | 5 | 4 |
| A | *Ceratotropis* | *Angulares* | *V. reflexopilosa* | 152 | 22.4 | 25.2 | 2439 | 1094 | 4 | 3 |
| A | *Ceratotropis* | *Angulares* | *V. reflexopilosa* ssp. *glabra* | 4 | na | na | na | na | 5 | 4 |
| A | *Ceratotropis* | *Angulares* | *V. riukiuensis* | 43 | 23.0 | 26.2 | 2374 | 834 | 3 | 1 |
| A | *Ceratotropis* | *Angulares* | *V. tenuicaulis* | 7 | 24.2 | 25.7 | 1267 | 670 | 5,5 | 2,3 |
| A | *Ceratotropis* | *Angulares* | *V. trinervia* | 37 | 23.9 | 23.6 | 2309 | 1022 | 5 | 3 |
| A | *Ceratotropis* | *Angulares* | *V. umbellata* | 627 | 22.1 | 24.9 | 1984 | 1007 | 4,5,5 | 3,4,4 |
| A | *Ceratotropis* | *Ceratotropis* | *V. grandiflora* | 4 | na | na | na | na | 5 | 3 |
| A | *Ceratotropis* | *Ceratotropis* | *V. hainiana* | 22 | 24.9 | 27.2 | 1248 | 926 | na | na |
| A | *Ceratotropis* | *Ceratotropis* | *V. mungo* | 891 | 21.3 | 25.1 | 997 | 603 | 4 | 2 |
| A | *Ceratotropis* | *Ceratotropis* | *V. mungo* var. *silvestris* | 16 | 24.9 | 25.5 | 1653 | 1009 | 5 | 3 |
| A | *Ceratotropis* | *Ceratotropis* | *V. radiata* | 1027 | 22.6 | 24.7 | 1085 | 573 | 5 | 4 |
| A | *Ceratotropis* | *Ceratotropis* | *V. radiata* var. *sublobata* | 548 | 25.4 | 27.5 | 1161 | 709 | 2,4 | 4,3 |
| A | *Ceratotropis* | *Ceratotropis* | *V. sahyadriana* | 0 | na | na | na | na | 5 | 4 |
| B | *Plectrotropis* | *Plectotropis* | *V. kirkii* | 77 | 23.9 | 24.3 | 1360 | 622 | na | na |
| B | *Plectrotropis* | *Plectotropis* | *V. vexillata* | 2394 | 21.8 | 23.4 | 1322 | 653 | 4,4,5,5 | 1,1,2,3 |
| B | *Plectrotropis* | *Plectotropis* | *V. vexillata* var. *angustifolia* | 940 | 22.3 | 24.9 | 1165 | 630 | 4 | 3 |
| B | *Plectrotropis* | *Plectotropis* | *V. vexillata* var. *davyi* | 56 | 17.3 | 20.3 | 991 | 486 | na | na |
| B | *Plectrotropis* | *Plectotropis* | *V. vexillata* var. *macrosperma* | 11 | 21.3 | 22.5 | 1439 | 683 | 4 | 1 |
| B | *Plectrotropis* | *Plectotropis* | *V. vexillata* var. *ovata* | 74 | 18.7 | 21.2 | 858 | 334 | 1 | 1 |
| B | *Plectrotropis* | *Plectotropis* | *V. vexillata* var. *youngiana* | 180 | 23.4 | 26.2 | 1181 | 689 | na | na |
| B | *Plectrotropis* | *Pseudoliebrechtsia* | *V. lobatifolia* | 1 | na | na | na | na | na | na |
| C | *Vigna* | *Catiang* | *V. keraudrenii* | 12 | 16.9 | 19.4 | 1404 | 827 | na | na |
| C | *Vigna* | *Catiang* | *V. monantha* | 8 | 26.3 | 26.6 | 184 | 87 | na | na |
| C | *Vigna* | *Catiang* | *V. schlechteri* | 136 | 16.1 | 19.3 | 974 | 509 | na | na |
| C | *Vigna* | *Catiang* | *V. unguiculata* | 6087 | 23.2 | 23.7 | 1172 | 626 | 4,4,4 | 1,2,2 |
| C | *Vigna* | *Catiang* | *V. unguiculata* group *biflora* | 23 | 20.4 | 23.8 | 1160 | 546 | na | na |
| C | *Vigna* | *Catiang* | *V. unguiculata* group *sesquipedalis* | 36 | 25.2 | 27.8 | 2850 | 1691 | 4 | 1 |
| C | *Vigna* | *Catiang* | *V. unguiculata* ssp. *aduensis* | 1 | na | na | na | na | na | na |
| C | *Vigna* | *Catiang* | *V. unguiculata* ssp. *alba* | 45 | 24.4 | 25.3 | 1378 | 571 | na | na |
| C | *Vigna* | *Catiang* | *V. unguiculata* ssp. *baoulensis* | 41 | 25.4 | 24.4 | 1352 | 661 | na | na |
| C | *Vigna* | *Catiang* | *V. unguiculata* ssp. *burundiensis* | 1 | na | na | na | na | na | na |
| C | *Vigna* | *Catiang* | *V. unguiculata* ssp. *dekindtiana* | 1016 | 23.0 | 24.2 | 936 | 508 | 3 | 2 |
| C | *Vigna* | *Catiang* | *V. unguiculata* ssp. *letouzeyi* | 21 | 25.1 | 24.3 | 1423 | 645 | na | na |
| C | *Vigna* | *Catiang* | *V. unguiculata* ssp. *pawekiae* | 45 | 20.3 | 21.0 | 1216 | 555 | na | na |
| C | *Vigna* | *Catiang* | *V. unguiculata* ssp. *protracta* | 112 | 18.4 | 21.6 | 879 | 392 | na | na |
| C | *Vigna* | *Catiang* | *V. unguiculata* ssp. *pubescens* | 58 | 25.1 | 25.8 | 1015 | 483 | na | na |
| C | *Vigna* | *Catiang* | *V. unguiculata* ssp. *stenophylla* | 228 | 19.6 | 22.5 | 816 | 396 | na | na |
| C | *Vigna* | *Catiang* | *V. unguiculata* ssp. *tenuis* | 108 | 20.8 | 23.2 | 945 | 500 | na | na |
| C | *Vigna* | *Catiang* | *V. unguiculata* ssp. *unguiculata* | 327 | 20.7 | 21.3 | 1023 | 532 | na | na |
| C | *Vigna* | *Catiang* | *V. unguiculata* var. *spontanea* | 58 | 23.0 | 24.5 | 873 | 469 | na | na |
| C | *Vigna* | *Catiang* | *V. unguiculata group melanophthalmus* | 0 | na | na | na | na | na | na |
| C | *Vigna* | *Macrodontae* | *V. friesiorum* | 41 | 19.8 | 20.2 | 951 | 415 | na | na |
| C | *Vigna* | *Macrodontae* | *V. frutescens* | 450 | 21.0 | 21.9 | 955 | 510 | na | na |
| C | *Vigna* | *Macrodontae* | *V. frutescens* ssp. *Incana* | 22 | 20.8 | 21.4 | 940 | 477 | na | na |
| C | *Vigna* | *Macrodontae* | *V. frutescens* var. *buchneri* | 32 | 20.2 | 20.9 | 1045 | 552 | na | na |
| C | *Vigna* | *Macrodontae* | *V. membranacea* | 301 | 21.6 | 21.6 | 978 | 467 | na | na |
| C | *Vigna* | *Macrodontae* | *V. somaliensis* | 0 | na | na | na | na | na | na |
| C | *Vigna* | *Reticulatae* | *V. reticulate* | 716 | 23.9 | 23.6 | 1233 | 651 | na | na |
| C | *Vigna* | *Reticulatae* | *V. wittei* | 30 | 22.6 | 21.7 | 1353 | 565 | na | na |
| D | *Vigna* | *Vigna* | *V. ambacensis* | 51 | 23.8 | 22.9 | 1252 | 614 | na | na |
| D | *Vigna* | *Vigna* | *V. angivensis* | 387 | 18.1 | 20.6 | 1413 | 841 | na | na |
| D | *Vigna* | *Vigna* | *V. bequaertii* | 29 | 19.3 | 19.2 | 1453 | 517 | na | na |
| D | *Vigna* | *Vigna* | *V. comosa* | 211 | 22.4 | 22.4 | 1522 | 650 | na | na |
| D | *Vigna* | *Vigna* | *V. filicaulis* | 101 | 26.5 | 25.5 | 1155 | 608 | na | na |
| D | *Vigna* | *Vigna* | *V. filicaulis* var. *pseudovenulosa* | 1 | na | na | na | na | na | na |
| D | *Vigna* | *Vigna* | *V. fischeri* | 2 | na | na | na | na | na | na |
| D | *Vigna* | *Vigna* | *V. gazensis* | 43 | 18.5 | 20.6 | 1235 | 715 | na | na |
| D | *Vigna* | *Vigna* | *V. gracilis* | 547 | 25.1 | 24.7 | 1653 | 794 | na | na |
| D | *Vigna* | *Vigna* | *V. heterophylla* | 512 | 25.0 | 24.4 | 1191 | 598 | na | na |
| D | *Vigna* | *Vigna* | *V. hosei* | 84 | 23.8 | 25.0 | 2153 | 894 | na | na |
| D | *Vigna* | *Vigna* | *V. laurentii* | 203 | 26.3 | 26.4 | 1196 | 561 | na | na |
| D | *Vigna* | *Vigna* | *V. luteola* | 1694 | 22.8 | 24.0 | 1422 | 627 | 4,5 | 1,1 |
| D | *Vigna* | *Vigna* | *V. marina* | 532 | 24.1 | 25.6 | 2123 | 870 | 4 | 1 |
| D | *Vigna* | *Vigna* | *V. multinervis* | 252 | 24.4 | 24.1 | 1403 | 661 | na | na |
| D | *Vigna* | *Vigna* | *V. oblongifolia* | 54 | 21.6 | 23.0 | 960 | 508 | na | na |
| D | *Vigna* | *Vigna* | *V. oblongifolia* var. *parviflora* | 304 | 21.3 | 22.3 | 951 | 465 | na | na |
| D | *Vigna* | *Vigna* | *V. o-wahuensis* | 151 | 19.9 | 18.8 | 1366 | 535 | na | na |
| D | *Vigna* | *Vigna* | *V. parkeri* | 90 | 19.3 | 20.2 | 1410 | 601 | na | na |
| D | *Vigna* | *Vigna* | *V. parkeri* ssp. *acutifolia* | 34 | 22.2 | 23.0 | 1254 | 623 | na | na |
| D | *Vigna* | *Vigna* | *V. parkeri* ssp. *maranguensis* | 177 | 19.7 | 19.8 | 1326 | 551 | na | na |
| D | *Vigna* | *Vigna* | *V. racemosa* | 751 | 25.7 | 24.9 | 1243 | 624 | na | na |
| D | *Vigna* | *Vigna* | *V. subterranea* | 1465 | 23.9 | 24.0 | 1069 | 618 | 4 | 2 |
| D | *Vigna* | *Vigna* | *V. subterranea* var. *spontanea* | 18 | 24.8 | 23.8 | 1013 | 592 | na | na |
| Color definition: **darkblue:** presence in harsh climate conditions, which are hot or dry compared to other taxa or high tolerance score in pot experiments; **intermediate blue**: presence in intermediate climates compared to other taxa or suboptimal tolerance score in pot experiments; **light blue**: presence in temperate climate conditions or humid climate conditions compared to other taxa or poor tolerance score in pot experiments. **Salmon**: no data. | | | | | | | | | | |
| ^I^ AMEANT: mean value of Annual Mean Temperature across taxa distribution; ^ii^ TWETQ: mean value of Temperature in the Wettest Quarter across taxa distribution; ^iii^ AP: mean value of Annual Precipitation across taxa distribution; ^iv^ PWETQ: mean value of Precipitation in the Wettest Quarter across taxa distribution; ^iv^ Drought tolerance score after lab screening carried out by Iseki et al. ^37^: 1 is high tolerance, 5 is low tolerance; ^v^ Salt tolerance score after lab screening carried out by Iseki et al. ^2^: 1 is high tolerance, 5 is low tolerance. | | | | | | | | | | |

**Table S3.** Biotic stress resistance of *Vigna* species

| Gene  pool | Subgenus | Section | Species | Insect and disease resistance found | References |
| --- | --- | --- | --- | --- | --- |
| A | *Ceratotropis* | *Aconitifoliae* | *V. aconitifolia* | YMD | ^38^ |
| A | *Ceratotropis* | *Aconitifoliae* | *V. subramaniana* | Bruchids (*C. chinensis*) | ^39^ |
| A | *Ceratotropis* | *Aconitifoliae* | *V. trilobata* | YMD | ^40^ |
| A | *Ceratotropis* | *Angulares* | *V. dalzelliana* | YMD | ^40^ |
| A | *Ceratotropis* | *Angulares* | *V. hirtella* | Bruchids (*Callosobruchus chinensis* and *C. macualtus*) | ^39^ |
| A | *Ceratotropis* | *Angulares* | *V. minima* | Bruchids (*C. chinensis* and *C. macualtus*) | ^41^ |
| A | *Ceratotropis* | *Angulares* | *V. nepalensis* | Bruchids (*C. chinensis* and *C. macualtus*) | ^39,40^ |
| A | *Ceratotropis* | *Angulares* | *V. reflexopilosa* | Bean fly (*Ophiomyia* phaseoli; *O. centrosematis*; *Melanagromyza sojae*); Bruchids (*Callosobruchus* spp.) | ^40,42^ |
| A | *Ceratotropis* | *Angulares* | *V. riukiuensis* | Bruchids (*Callosobruchus* spp.) | ^42^ |
| A | *Ceratotropis* | *Angulares* | *V. tenuicaulis* | Bruchids (*Callosobruchus* spp.) | ^41^ |
| A | *Ceratotropis* | *Angulares* | *V. trinervia* | YMD | ^40^ |
| A | *Ceratotropis* | *Angulares* | *V. umbellata* | Bruchids (*Callosobruchus* spp.); YMD | ^40,41^ |
| A | *Ceratotropis* | *Angulares* | *V. reflexopilosa* ssp. *glabra* | Bruchids (*Callosobruchus* spp.); bean fly; YMD; CMV; Powdery mildew | ^12,40^ |
| A | *Ceratotropis* | *Angulares* | *V. trinervia* | Bruchids (*C. chinensis*) | ^39^ |
| A | *Ceratotropis* | *Ceratotropis* | *V. radiata* | Anthracnose (*Colletotrichum lindemuthianum* or *C. truncatum* or *C. gloeosporioides*); Bean blossom thrips (*Megalurothrips distalis)*; Cotton bollworm (*Helicoverpa armigera*); Cowpea aphid (*Aphis craccivora*); Cercospora leaf spot (*Cercospora cruenta* or *C. canescens* or *C. kikuchii* or *C. caracallae*); Dry root rot (*Rhizoctonia bataticola*); Green Jassid (*Empoasca* spp.); Legume pod borer (*Maruca vitrata*); Macrophomina blight (*M. phaseolina*); Powdery mildew (*Erysiphe polygoni* or *Podosphaera fusca*); Stem borer (*Ophiomyia* spp.); Whitefly (*Bemisia tabaci*); YMD | ^40,43,44^ |
| A | *Ceratotropis* | *Ceratotropis* | *V. mungo* | Bean blossom thrips (*M. distalis*); Bruchids (*C. chinensis*); Cotton bollworm (*H. armigera*); Cowpea aphid (*A. craccivora*); Whitefly (*B. tabaci*); YMD | ^44,45^ |
| A | *Ceratotropis* | *Ceratotropis* | *V. mungo* var. *silvestris* | Bruchids (*Callosobruchus* spp.) | ^40^ |
| A | *Ceratotropis* | *Ceratotropis* | *V. radiata* var. *sublobata* | Bruchids (*Callosobruchus* spp.); YMD | ^40^ |
| B | *Plectrotropis* | *Plectotropis* | *V. vexillata* | Bruchids (*C. maculatus*); CPMoV | ^40^ |
| C | *Vigna* | *Catiang* | *V. unguiculata* | Anthracnose (*Colletotrichum destructivum*); Bacterial blight (*Xanthomonas vignicola* Burkh.); Bean Bug (*Clavigralla tomentosicollis*); Bean fly (*O. phaseoli*); BICMV; Bruchids (*C. maculatus*); CAMV; Cowpea aphid (*A. craccivora*); Cowpea flower thrips (*M. sjostedti*); Leafhoppers (*Empoasca* spp.); Legume pod borer (*M. vitrata);* Root‑knot nematodes *(Meloidogyne* spp.); Thrips (*Frankliniella* spp.); YMD | ^45–48^ |
| C | *Vigna* | *Catiang* | *V. unguiculata* ssp. *mensensis* | Cowpea moth (*Cydia ptychor*) | ^40^ |
| C | *Vigna* | *Catiang* | *V. unguiculata* ssp. *dekindtiana* | Coreid bug (*Clavigralla tomentosicollis*) | ^40^ |
| C | *Vigna* | *Reticulatae* | *V. reticulata* | Bruchids (*C. maculatus*) | ^49^ |
| D | *Vigna* | *Vigna* | *V. luteola* | Bruchids (*C. maculatus*) | ^49^ |
| D | *Vigna* | *Vigna* | *V. oblongifolia* | Bruchds (*C. maculatus*) | ^49^ |
| CAMV: Cowpea aphid-borne mosaic virus; BICMV: Blackeye cowpea mosaic virus; CPMoV: Cowpea mottle carmovirus; YMD: Yellow mosaic disease viruses. | | | | | |

**Table S4.** Reported accessions conserved *ex situ* in 2017

| Gene  pool | Subgenus | Section | | Species | Meise, Belgium | | IITA | | Australian Grains Genebank | | CIAT | | NARO, Japan | | NBPGR, India | | WorldVeg | | Subtotal | | Total | |
| --- | --- | --- | --- | --- | --- | --- | --- | --- | --- | --- | --- | --- | --- | --- | --- | --- | --- | --- | --- | --- | --- | --- |
| A | *Ceratotropis* | *Angulares* | | *V. angularis* | 9 | |  | | 349 | | 3 | | 1492 | | 175 | | 2350 | | 4378 | | 5082 | |
| A | *Ceratotropis* | *Angulares* | | *V. dalzelliana* |  | |  | | 4 | | 4 | |  | | 21 | |  | | 29 | | 36 | |
| A | *Ceratotropis* | *Angulares* | | *V. exilis* | 1 | |  | |  | |  | |  | |  | |  | | 1 | | 1 | |
| A | *Ceratotropis* | *Angulares* | | *V. hirtella* | 3 | |  | |  | |  | | 4 | |  | |  | | 7 | | 7 | |
| A | *Ceratotropis* | *Angulares* | | *V. minima* | 2 | | 1 | |  | | 2 | | 5 | | 1 | |  | | 11 | | 12 | |
| A | *Ceratotropis* | *Angulares* | | *V. nakashimae* | 2 | |  | |  | |  | | 21 | |  | |  | | 23 | | 24 | |
| A | *Ceratotropis* | *Angulares* | | *V. nepalensis* | 3 | |  | |  | |  | | 4 | | 3 | |  | | 10 | | 10 | |
| A | *Ceratotropis* | *Angulares* | | *V. reflexopilosa* | 3 | | 2 | | 1 | | 2 | | 37 | |  | | 3 | | 48 | | 50 | |
| A | *Ceratotropis* | *Angulares* | | *V. riukiuensis* | 1 | |  | |  | |  | | 63 | |  | |  | | 64 | | 64 | |
| A | *Ceratotropis* | *Angulares* | | *V. tenuicaulis* | 1 | |  | |  | |  | | 2 | |  | |  | | 3 | | 3 | |
| A | *Ceratotropis* | *Angulares* | | *V. trinervia* | 2 | |  | |  | |  | | 6 | | 8 | |  | | 16 | | 16 | |
| A | *Ceratotropis* | *Angulares* | | *V. umbellata* | 13 | | 1 | | 59 | | 39 | | 214 | | 2050 | | 320 | | 2696 | | 3003 | |
| A | *Ceratotropis* | *Ceratotropis* | | *V. grandiflora* | 1 | |  | |  | |  | | 1 | |  | |  | | 2 | | 2 | |
| A | *Ceratotropis* | *Ceratotropis* | | *V. hainiana* |  | |  | |  | |  | |  | | 2 | |  | | 2 | | 2 | |
| A | *Ceratotropis* | *Ceratotropis* | | *V. mungo* | 12 | | 11 | | 104 | | 96 | | 145 | | 1751 | | 849 | | 2968 | | 5987 | |
| A | *Ceratotropis* | *Ceratotropis* | | *V. radiata* | 34 | | 124 | | 1385 | | 69 | | 922 | | 4024 | | 6752 | | 13310 | | 21161 | |
| A | *Ceratotropis* | *Ceratotropis* | | *V. sahyadriana* |  | |  | |  | |  | |  | |  | |  | | 0 | | 0 | |
| A | *Ceratotropis* | *Aconitifoliae* | | *V. aconitifolia* | 7 | |  | | 35 | | 8 | | 6 | | 1486 | | 26 | | 1568 | | 1849 | |
| A | *Ceratotropis* | *Aconitifoliae* | | *V. aridicola* | 1 | |  | |  | |  | |  | |  | |  | | 1 | | 1 | |
| A | *Ceratotropis* | *Aconitifoliae* | | *V. indica* |  | |  | |  | |  | |  | |  | |  | | 0 | | 0 | |
| A | *Ceratotropis* | *Aconitifoliae* | | *V. khandalensis* |  | |  | |  | |  | |  | | 1 | |  | | 1 | | 1 | |
| A | *Ceratotropis* | *Aconitifoliae* | | *V. stipulacea* | 4 | |  | |  | |  | | 1 | |  | |  | | 5 | | 5 | |
| A | *Ceratotropis* | *Aconitifoliae* | | *V. subramaniana* | 1 | |  | | 2 | |  | | 1 | |  | |  | | 4 | | 6 | |
| A | *Ceratotropis* | *Aconitifoliae* | | *V. trilobata* | 4 | | 7 | | 53 | | 4 | |  | | 130 | | 2 | | 200 | | 336 | |
| B | *Plectrotropis* | *Plectotropis* | | *V. kirkii* | 1 | | 6 | | 1 | |  | |  | |  | |  | | 8 | | 9 | |
| B | *Plectrotropis* | *Plectotropis* | | *V. vexillata* | 135 | | 195 | | 187 | | 201 | | 6 | | 110 | | 2 | | 836 | | 1068 | |
| B | *Plectrotropis* | *Pseudoliebrechtsia* | | *V. lobatifolia* |  | | 3 | | 1 | |  | |  | |  | |  | | 4 | | 5 | |
| C | *Vigna* | *Catiang* | | *V. keraudrenii* |  | |  | |  | |  | |  | |  | |  | | 0 | | 0 | |
| C | *Vigna* | *Catiang* | | *V. monantha* |  | |  | |  | |  | |  | |  | |  | | 0 | | 0 | |
| C | *Vigna* | *Catiang* | | *V. slechteri* |  | | 9 | |  | |  | |  | |  | |  | | 9 | | 9 | |
| C | *Vigna* | *Catiang* | | *V. unguiculata* | 332 | | 16127 | | 935 | | 94 | | 1371 | | 3649 | | 1610 | | 24118 | | 44694 | |
| C | *Vigna* | *Macrodontae* | | *V. friesiorum* | 1 | | 6 | | 1 | | 1 | |  | |  | |  | | 9 | | 11 | |
| C | *Vigna* | *Macrodontae* | | *V. frutescens* | 14 | | 7 | | 10 | | 3 | |  | |  | |  | | 34 | | 46 | |
| C | *Vigna* | *Macrodontae* | | *V. membranacea* | 32 | | 13 | | 3 | | 1 | |  | |  | |  | | 49 | | 96 | |
| C | *Vigna* | *Macrodontae* | | *V. somaliensis* |  | |  | |  | |  | |  | |  | |  | | 0 | | 0 | |
| C | *Vigna* | *Reticulatae* | | *V. reticulata* | 28 | | 116 | | 3 | |  | |  | |  | |  | | 147 | | 202 | |
| C | *Vigna* | *Reticulatae* | | *V. wittei* |  | | 29 | |  | | 1 | |  | |  | |  | | 30 | | 30 | |
| D | *Vigna* | *Vigna* | | *V. ambacensis* | 36 | | 151 | | 4 | | 1 | |  | |  | |  | | 192 | | 195 | |
| D | *Vigna* | *Vigna* | | *V. angivensis* | 3 | | 1 | | 1 | |  | |  | |  | |  | | 5 | | 6 | |
| D | *Vigna* | *Vigna* | | *V. bequaertii* |  | |  | |  | |  | |  | |  | |  | | 0 | | 0 | |
| D | *Vigna* | *Vigna* | | *V. comosa* | 1 | | 11 | |  | |  | |  | |  | |  | | 12 | | 13 | |
| D | *Vigna* | *Vigna* | | *V. filicaulis* | 5 | | 4 | | 2 | |  | |  | |  | |  | | 11 | | 14 | |
| D | *Vigna* | *Vigna* | | *V. fischeri* |  | | 1 | |  | |  | |  | |  | |  | | 1 | | 1 | |
| D | *Vigna* | *Vigna* | | *V. gazensis* |  | |  | |  | |  | |  | |  | |  | | 0 | | 0 | |
| D | *Vigna* | *Vigna* | | *V. gracilis* | 14 | | 24 | |  | |  | |  | |  | |  | | 38 | | 49 | |
| D | *Vigna* | *Vigna* | | *V. heterophylla* | 2 | | 1 | | 4 | |  | |  | |  | |  | | 7 | | 32 | |
| D | *Vigna* | *Vigna* | | *V. hosei* | 5 | | 44 | | 4 | | 7 | |  | |  | |  | | 60 | | 65 | |
| D | *Vigna* | *Vigna* | | *V. laurentii* | 2 | | 1 | | 1 | |  | |  | |  | |  | | 4 | | 6 | |
| D | *Vigna* | *Vigna* | | *V. luteola* | 24 | | 66 | | 39 | | 69 | |  | |  | | 4 | | 202 | | 257 | |
| D | *Vigna* | *Vigna* | | *V. marina* | 6 | | 8 | | 39 | |  | | 1 | | 2 | | 43 | | 99 | | 129 | |
| D | *Vigna* | *Vigna* | | *V. multinervis* | 5 | | 17 | |  | |  | |  | |  | |  | | 22 | | 23 | |
| D | *Vigna* | *Vigna* | | *V. oblongifolia* | 31 | | 55 | | 42 | | 31 | |  | |  | |  | | 159 | | 256 | |
| D | *Vigna* | *Vigna* | | *V. o-wahuensis* | 1 | |  | |  | |  | |  | |  | |  | | 1 | | 1 | |
| D | *Vigna* | *Vigna* | | *V. parkeri* | 6 | | 2 | | 20 | | 1 | |  | |  | | 1 | | 30 | | 80 | |
| D | *Vigna* | *Vigna* | | *V. racemosa* | 43 | | 131 | | 6 | | 1 | |  | |  | |  | | 181 | | 208 | |
| D | *Vigna* | *Vigna* | | *V. subterranea* |  | | 2088 | | 47 | |  | |  | |  | |  | | 2135 | | 4125 | |
|  |  | | Number of accessions | | | 831 | | 19262 | | 3342 | | 638 | | 4302 | | 13413 | | 11962 | | 53750 | | 89288 |
|  |  | | Number of taxa | | | 41 | | 32 | | 29 | | 21 | | 19 | | 15 | | 12 | | 55 | | 55 |

**Table S5.** Targeted countries for collecting of *V.* species with less than 10 genebank accessions

| Subgenus | Section | Taxa | Reported occurrence | Modelled occurrence |
| --- | --- | --- | --- | --- |
| *Ceratotropis* | *Angulares* | *V. exilis* | **Thailand ^i^** | **Thailand**, Myanmar |
| *Ceratotropis* | *Angulares* | *V. hirtella* | **Thailand**, India, Myanmar, Malaysia, Vietnam, China, Lao People's Democratic Republic | **Myanmar**, Thailand, India, Cambodia, China, Malaysia, Indonesia, Bhutan, Nepal, Bangladesh |
| *Ceratotropis* | *Angulares* | *V. tenuicaulis* | **Thailand** | **Thailand** |
| *Ceratotropis* | *Ceratotropis* | *V. grandiflora* | **Thailand**, Cambodia | **Thailand**, Cambodia |
| *Ceratotropis* | *Ceratotropis* | *V. hainiana* | **India** | **India**, Nepal |
| *Ceratotropis* | *Ceratotropis* | *V. sahyadriana* | na | na |
| *Ceratotropis* | *Aconitifoliae* | *V. aridicola* | **Sri Lanka** | **Sri Lanka** |
| *Ceratotropis* | *Aconitifoliae* | *V. indica* | na | na |
| *Ceratotropis* | *Aconitifoliae* | *V. khandalensis* | **India** | **India** |
| *Ceratotropis* | *Aconitifoliae* | *V. stipulacea* | **India**, Indonesia, Sri Lanka, Spain, Vietnam | **India**, Thailand, Myanmar, Indonesia, Cambodia, Australia, Philippines, Sri Lanka, Ethiopia, Bangladesh, South Sudan, Malaysia, Sudan, China, Maldives |
| *Ceratotropis* | *Aconitifoliae* | *V. subramaniana* | **India** | **India**, Pakistan, Nepal, China |
| *Plectrotropis* | *Plectotropis* | *V. kirkii* | **Tanzania**, DRC Congo, Uganda, Mozambique, Malawi, Burundi, Cameroon, Guinea-Bissau, Guinea, Kenya, Senegal, South Sudan, Zambia | **DRC Congo**, Central African Republic, Mozambique, Cameroon, Congo, Nigeria, Uganda, Ghana, South Sudan, Angola, Gabon, Malawi, Kenya, Zambia, Guinea, Ethiopia, Senegal, Guinea-Bissau, Mali, Togo, Benin, Liberia, Burundi, Burkina Faso, Rwanda, Madagascar, Chad, Equatorial Guinea, Zimbabwe, Sierra Leone, Gambia, Comoros |
| *Plectrotropis* | *Pseudoliebrechtsia* | *V. lobatifolia* | **Namibia** | na |
| *Vigna* | *Catiang* | *V. keraudrenii* | **Madagascar** | **Madagascar** |
| *Vigna* | *Catiang* | *V. monantha* | **Somalia** | **Somalia** |
| *Vigna* | *Catiang* | *V. schlechteri* | **South Africa**, Zimbabwe, Swaziland, Mozambique | South Africa, Zimbabwe, Swaziland, Mozambique, Lesotho |
| *Vigna* | *Macrodontae* | *V. somaliensis* | na | na |
| *Vigna* | *Vigna* | *V. angivensis* | **Madagascar**, Burundi, Russian Federation | **Madagascar**, Nepal, India, Mozambique, Democratic Republic of the Congo, Reunion (France), Comoros, Mauritius, Zimbabwe, Kenya, China, Uganda, Malawi, Ethiopia, Rwanda |
| *Vigna* | *Vigna* | *V. bequaertii* | **DRC Congo**, Burundi, Rwanda | **DRC Congo**, Central African Republic, Rwanda, Uganda, Burundi, South Sudan, Angola |
| *Vigna* | *Vigna* | *V. fischeri* | **Burundi**, Malawi | na |
| *Vigna* | *Vigna* | *V. gazensis* | **Mozambique**, Malawi, Madagascar, Zimbabwe | **Madagascar**, Mozambique, Zimbabwe, Malawi, Zambia |
| *Vigna* | *Vigna* | *V. laurentii* | **Benin**, DRC Congo, Cameroon, Burundi, Togo, Central African Republic, Gabon, Guinea-Bissau, South Sudan | **DRC Congo**, Nigeria, Central African Republic, Cameroon, Angola, South Sudan, Zambia, Ghana, Gabon, Congo, Uganda, Chad, Benin, Togo, Guinea, Burundi, Guinea-Bissau, Rwanda, Burkina Faso, Senegal, Mali, Equatorial Guinea, Sudan, Kenya, Sao Tome and Principe, Malawi, Sierra Leone |
| *Vigna* | *Vigna* | *V. o-wahuensis* | **United States** | **United States**, Canada, Mexico |
| ^i^ Countries are presented in order of occurrence**. Countries in bold** reported most records of the corresponding species and most coverage of its modelled distribution | | | | |
